# Matching whole genomes to rare genetic disorders: Identification of potential causative variants using phenotype-weighted knowledge in the CAGI SickKids5 clinical genomes challenge

**DOI:** 10.1101/707687

**Authors:** Lipika R. Pal, Kunal Kundu, Yizhou Yin, John Moult

## Abstract

Precise identification of causative variants from whole-genome sequencing data, including both coding and non-coding variants, is challenging. The CAGI5 SickKids clinical genome challenge provided an opportunity to assess our ability to extract such information. Participants in the challenge were required to match each of 24 whole-genome sequences to the correct phenotypic profile and to identify the disease class of each genome. These are all rare disease cases that have resisted genetic diagnosis in a state-of-the-art pipeline. The patients have a range of eye, neurological, and connective-tissue disorders. We used a gene-centric approach to address this problem, assigning each gene a multi-phenotype-matching score. Mutations in the top scoring genes for each phenotype profile were ranked on a six-point scale of pathogenicity probability, resulting in an approximately equal number of top ranked coding and non-coding candidate variants overall. We were able to assign the correct disease class for 12 cases and the correct genome to a clinical profile for five cases. The challenge assessor found genes in three of these five cases as likely appropriate. In the post-submission phase, after careful screening of the genes in the correct genome we identified additional potential diagnostic variants, a high proportion of which are non-coding.

## INTRODUCTION

Identification of the variant(s) causing a patient’s clinical symptoms is one of the key challenges in rare disease diagnostics. The problem has assumed increasing urgency, with recent advances in sequencing technology and a decrease in the sequencing cost (Schwarze, Buchanan, Taylor, & Wordsworth, 2018), leading to vast amounts of data to interpret. While whole-genome sequencing data provides more comprehensive coverage than other more restricted sequencing technologies (such as gene panel data, exome sequencing data, or chromosomal microarray data), identification of potential causative variant(s) out of the approximately four million variants found in a genome resonates with finding a needle in a haystack (Cooper & Shendure, 2011). The variant diagnostic rate (rate of causative, pathogenic or likely pathogenic genotypes in known disease genes for children) from whole-genome sequencing data is currently only about 40% (Clark et al., 2018) with the implication that there remain substantial deficiencies in current methodology. Many factors contribute to this shortfall, but there is a clear need for improved methods from the computational biology community. The Critical Assessment of Genome Interpretation (CAGI) (https://genomeinterpretation.org) is a platform for community experiments in genome interpretation. Typically, the experiments take the form of blinded prediction of the phenotypic impacts of genomic variation followed by an objective independent assessment of the results (Hoskins et al., 2017). The SickKids5 experiment is one such challenge (https://genomeinterpretation.org/SickKids5_clinical_genomes), and follows an earlier CAGI SickKids4 one (https://genomeinterpretation.org/content/4-SickKids_clinical_genomes). Here we report our methods and the results obtained for this challenge, and draw conclusions on directions for future improvement.

Participants were provided with a set of 24 whole genome sequencing data and 24 clinical profiles for pediatric patients, and asked to match each genome to the corresponding phenotype profile. Two general strategies have been developed: genotype to phenotype and phenotype to genotype. In a genotype to phenotype approach, a patient’s clinical profile is not utilized to prioritize potential causative genes and variant(s)-rather all deleterious variants in a genome are identified from genotype data. Several variant annotation programs (such as VAAST (Hu et al., 2013), ANNOVAR (Wang, Li, & Hakonarson, 2010), SnpEff (Cingolani, Platts, et al., 2012), VAT (Habegger et al., 2012), and VEP (McLaren et al., 2010)) utilize population allele frequency data and evolutionary conservation information together with appropriate disease inheritance models to prioritize disease relevant genes and variants in a genome, without explicitly considering a specific patient phenotypic profile. Conversely, the common theme of a phenotype to genotype approach is that a set of patient specific phenotypes, either in the form of Human Phenotype Ontology (HPO) (Köhler et al., 2014) terms or other clinical descriptors, is used to generate a list of relevant genes and only variants in these genes are considered for further analysis. A number of strategies have been developed for incorporation of patient-specific gene prioritization information. The information may come from various bio-medical ontologies, including human-specific ontologies, like HPO (Köhler et al., 2014), DO (Schriml et al., 2019), GO (Blake et al., 2015) (such as used in Phevor (Singleton et al., 2014)) and other model organism specific ontologies, such as MPO (Smith & Eppig, 2009), ZPO (van Slyke, Bradford, Westerfield, & Haendel, 2014) (used in Exomiser (Smedley et al., 2015)). Several computational tools leverage gene-disease-phenotype relationships and phenotype information, for instance Phenolyzer (H. Yang, Robinson, & Wang, 2015), and PDR (Krämer, Shah, Rebres, Tang, & Richards, 2017). Also some tools extract phenotype information using keyword search from text (used in GeneCards (Safran et al., 2010)), or free-text boolean search (used in VarElect (Stelzer et al., 2016)). Prioritization of variants in prioritized genes includes evaluation of the likely impact on gene function and pruning variants on the basis of population frequency data (rare disease implies rare variants). Computational tools use a variety of methodologies to assess the likely impact of coding and non-coding variants. Evaluation of coding variants is usually divided into Loss of Function (frameshifts, direct splice site impact, and stop gain or loss) and missense. Many methods have been developed to estimate the effect of missense mutations based on sequence conservation properties (for example, SIFT (Kumar, Henikoff, & Ng, 2009), PolyPhen-2 (Adzhubei et al., 2010), SNPs3D profile (Yue & Moult, 2006), SNAP2 (Hecht, Bromberg, & Rost, 2015), and Evolutionary Action (Katsonis & Lichtarge, 2017)) and on protein stability, as estimated from protein structure (for example, SNPs3D stability (Yue, Li, & Moult, 2005), Rosetta (Park et al., 2016), and FoldX (Delgado, Radusky, Cianferoni, & Serrano, 2019; Schymkowitz et al., 2005)). Some methods also include functional information (for example, MutPred2 (Pejaver, Mooney, & Radivojac, 2017)). Non-coding variant analysis methods utilize features including regional purifying selection, enrichment with functional elements such as transcription factor binding sites and DNase hypersensitivity as well as DNA based evolutionary conservation. Example methods are PhastCons (Siepel et al., 2005), PhyloP (Pollard, Hubisz, Rosenbloom, & Siepel, 2010), and Gerp++ (Davydov et al., 2010). Features are often combined using machine learning approaches (such as those used in Genomiser (Smedley et al., 2016) and CADD (Rentzsch, Witten, Cooper, Shendure, & Kircher, 2019)). In one analysis (Smedley & Robinson, 2015), phenotype-driven approaches were found to have substantially better performance than variant driven ones. So far, most of these methodologies have only been benchmarked against simulated data, and there has been very limited blind testing.

In the Sickkids5 challenge, participants were provided with clinical profiles in the form of a set of PhenoTips terms (Girdea et al., 2013) (represented using HPO terms) and whole genome sequencing data for the 24 pediatric patients. These are all difficult cases where the standard SickKids analysis pipeline failed to find any reportable diagnostic variants (Kasak et al., 2019). The challenge was to assign each genome to one of three disease classes (eye disorders, neurological disorders, and connective tissue disorders) and to match each genome to the appropriate clinical profile. An additional optional part to the challenge was the identification of specific diagnostic variants for each patient. The identification of predictive secondary variants (related to risk of other serious diseases and with no phenotypes reported in the clinical descriptions) was also optional.

Here we report our approach and results for the SickKids5 challenge. We used a phenotype to genotype approach, selecting only clinical symptom-specific genes. For this purpose, we developed a phenotype-weighted scoring scheme to select the set of genes associated with each clinical profile. Each variant in the selected genes was assigned to one of six impact related categories. Final selection of a genome for each clinical profile included a subjective evaluation of the match of each gene’s OMIM description (Hamosh, Scott, Amberger, Bocchini, & McKusick, 2005) with the clinical profile. The results were analyzed in a number of ways, especially the role of clear clinical documentation in developing the phenotype-weighted scoring scheme and types of prioritized variants.

## MATERIALS AND METHODS

### SickKids5 clinical profile data

The SickKids Genome Clinic at the Hospital for Sick Children in Toronto (http://www.sickkids.ca/) provided the clinical profiles for the challenge. The profiles for the 24 patients included an overall disease class, with six eye disorder cases, seven neurological and 11 connective-tissue disorders cases. Additional profile information for each patient included gender, age, indication for referral and clinical symptoms in the form of a set of terms from the hierarchical Human Phenotype Ontology (HPO) (Köhler et al., 2014) entered through the PhenoTips interface (Girdea et al., 2013). Inheritance information was also provided for some patients: Six were described as autosomal recessive cases and pedigree charts were given for 14 patients (including two of the six autosomal recessive cases). Ethnicity information was also provided for 19 out of 24 patients, none of whom were declared as African origin. We used in-house software to identify genomes of African origin (described in (Pal, Kundu, Yin, & Moult, 2017)). In the post-challenge submission phase, using the answer key, we found that one patient with declared Philippine ethnicity is genetically of African origin, and this caused a prediction error.

### SickKids5 whole genome data: annotation of VCF files and QC filters

Anonymized whole genome data for all 24 patients were available via the CAGI SickKids5 challenge website (https://genomeinterpretation.org/SickKids5_clinical_genomes) in the form of VCF files produced by the Illumina HiSeq X system. We annotated SNVs and Indels in the VCF files using Varant (https://doi.org/10.5060/D2F47M2C), including region of occurrence (intron, exon, splice site, or intergenic), observed minor allele frequencies (MAF), mutation type, predicted impact on protein function (methods used in this step are listed in ‘Categorization of variants’ section under Methods), and associated phenotypes reported in ClinVar (Landrum et al., 2016). The RefGene (Pruitt et al., 2014) gene definition file was used for gene and transcript annotations in Varant. In addition, in-house scripts were used to annotate variants with HGMD (Stenson et al., 2014) disease related information and with dbscSNV (Jian, Boerwinkle, & Liu, 2014) information on potential splicing effects. We also used Annovar annotations (Wang et al., 2010) to add Genome Aggregation Database (GnomAD) frequency data (Lek et al., 2016), Eigen scores (Ionita-Laza, McCallum, Xu, & Buxbaum, 2016) and GERP++ scores (Davydov et al., 2010) information for each variant. Chromosome M was annotated and searched for pathogenic variants using MSeqDR mv (Shen et al., 2018). We used only high quality (graded ‘PASS’ in the VCF file) variants for further analysis. We used SnpSift (Cingolani, Patel, et al., 2012) to calculate Ts/Tv and Het/Hom alternate allele ratios from the VCF file data. We only considered variants for which the highest population frequency is <1% in all the referenced databases (GnomAD exomes and GnomAD genomes, ExAC database (Lek et al., 2016), and 1000 genomes (Auton et al., 2015)).

### Method rationale

In order to address the challenge of matching genomes to clinical profiles and identifying the disease class of each genome, we used a phenotype to genotype approach, first identifying genes compatible with clinical profile information, and then analyzing variants in those genes. If we are able to identify an appropriate candidate causal variant (or pair if variants for a recessive trait) for a specific profile, that is taken as evidence of a genome and profile match, and will also imply the disease class of that genome. The steps in the method are: (1) Collection of disease relevant genes for a particular clinical profile from all 24 genomes (details in ‘Candidate gene list generation’ section under Methods); (2) Identification of rare variants (less than 1% population frequency) in the relevant genes (as mentioned earlier in ‘SickKids5 whole genome data’ section); (3) Search for impact variants (both coding and non-coding) in the relevant genes and assignment of these to one of the six categories of impact confidence (details in ‘Categorization of variants’ section); (4) Use of a subjective scoring scheme of the clinical profile (depending on the presumed disease class to which that particular profile belongs) to score each disease relevant gene in a genome for each clinical profile (details in ‘Gene scoring scheme’); (5) selection of the top five scoring genomes for each clinical profile, and within those, selection of top five scoring genes; and (6) manual screening of the variants selected for each profile for appropriate inheritance model, ethnicity compatibility, and the full match of the OMIM disease description associated with each gene to the clinical profile (details in ‘Prioritized causative variants for a genome’ section). The work flow of the method is shown in Figure 1.

**Figure 1:**
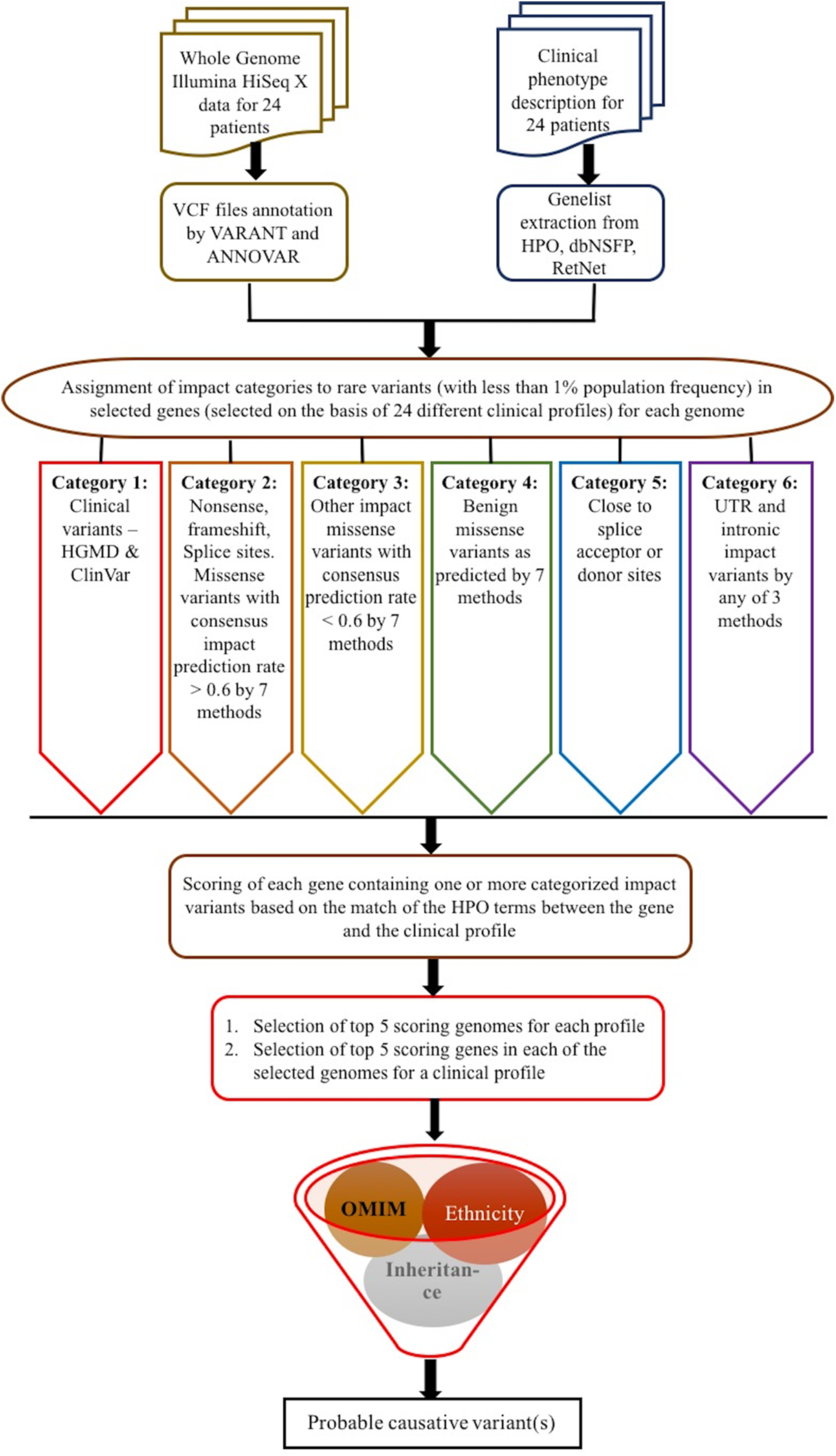
Workflow of the method for identification of probable causative variants.

### Candidate gene list generation

For each patient, we extracted the Human Phenotype Ontology-based (HPO) terms from the PhenoTips annotations provided in the clinical profile. Relevant genes for each profile were identified by matching the profile HPO terms to those associated with each gene in the HPO database (Build #139) (Köhler et al., 2014) and in the dbNSFP database (version 3.5a) (Liu, Wu, Li, & Boerwinkle, 2016). The latter includes genes related to phenotypes observed in humans as well as similar phenotypes included in the mouse database (Eppig et al., 2015; Georgi, Voight, & Bućan, 2013). We also used the list of 319 genes from the RetNet database (RetNet, http://www.sph.uth.tmc.edu/RetNet/) (Daiger, 2004) to search for eye disorder related variants. The gene list for secondary variants, containing 59 genes, was taken from the Table 1 in the 2017 ACMG guidelines (Kalia et al., 2017).

**Table 1:** Performance of the method in CAGI SickKids5 challenge.

| **Broad disease class** | **Total number of cases** | **Number of cases with correct disease class assignment** | **Number of cases with exact match between clinical profile and genome** | **Number of cases with proposed diagnostic genes considered probable by the assessor** |
| --- | --- | --- | --- | --- |
| Connective tissue disorder | 11 | 6 | 2 | 1 |
| Eye disorder | 6 | 3 | 2 | 2 |
| Neurological disorder | 7 | 3 | 1 | 0 |

### Categorization of variants according to their likely pathogenic impact

As outlined in the rationale, for each genome we identified 24 different sets of possible candidate causative variants, one for each of the clinical profiles. Only variants with less than 1% population frequency were considered. Each selected variant was assigned to one of six categories, based on likelihood of pathogenicity and variant type, as follows: Category 1 (C1): Variants with HGMD annotation of either DM (disease-causing mutation) or DP (disease-associated polymorphism), and/or reported in ClinVar with pathogenic or likely pathogenic clinical significance status.

Category 2 (C2): Nonsense mutation, frameshift or non-frameshift indel, a mutation disrupting either a splice donor or acceptor site, splice altering variants (splicing consensus regions around direct splice sites) predicted by the dbscSNV (Jian et al., 2014), and missense mutations predicted as damaging by SNPs3D profile and stability methods (Yue, Li, & Moult, 2005; Yue & Moult, 2006), SIFT (Kumar, Henikoff, & Ng, 2009), PolyPhen-2 (Adzhubei et al., 2010), Vest (Carter, Douville, Stenson, Cooper, & Karchin, 2013), REVEL (Ioannidis et al., 2016) and CADD (Kircher et al., 2014). For inclusion of a missense mutation in Category 2, at least 60% of reporting methods were required to return a prediction of deleterious. This threshold is based on a calibration against HGMD (Yin, Kundu, Pal, & Moult, 2017).

Category 3 (C3): Missense mutations predicted as damaging by one or more of the above missense impact prediction methods, with the fraction of deleterious predictions < 0.6.

Category 4 (C4): Benign missense mutations (zero reporting missense methods predicting deleterious).

Category 5 (C5): Variants annotated as close (within 12 bases) to a splice acceptor or splice donor site.

Category 6 (C6): Variants annotated as UTR and intronic with at least one of the following conditions satisfied: CADD phred score > 20 (Kircher et al., 2014), Eigen score >=4 (Ionita-Laza et al., 2016), or Gerp++ score >=2 (Davydov et al., 2010).

Variants in all categories were further subdivided on the basis of population frequency data: Frequency bin 1: Novel mutations (not seen in any of 1000 genomes, ExAC, gnomAD exomes and gnomAD genomes databases).

Frequency bin 2: Variants with population frequency > 0 and <= 0.001.

Frequency bin 3: Variants with population frequency > 0.001 and <= 0.005.

Frequency bin 4: Variants with population frequency > 0.005 and < 0.01.

Variants were assigned to autosomal dominant, autosomal recessive, compound heterozygous, pseudo autosomal recessive, or X-linked recessive models based on the OMIM inheritance pattern for the corresponding gene (https://www.ncbi.nlm.nih.gov/omim).

The subset of selected genes in a genome that contain one or more impact variants are then considered in scoring of genome’s match to a clinical profile.

### Gene scoring scheme for selection of genomes best matching to a clinical profile

For each clinical profile, each HPO term (T) was assigned a subjective weight (W) from 0 to 1, according to its importance (1 = most important and 0 = least important) in that profile, taking into account the presumed disease class. Usually the most important terms were inferred from the ‘indication for referral’ description. For example, if a connective tissue disorder (presumed disease class of that clinical profile) is the most dominant and definitive term in the profile in the ‘indication for referral’ description, it was scored the highest. If seizure is also part of that profile but with borderline occurrence, then that was assigned a lower value than would be the case if the term occurred in a profile where seizure is the most significant phenotype in the ‘indication for referral’ field.

We started with the set of genes containing impact variants identified in each genome. For each clinical profile, each selected gene ‘i’ of a genome was assigned a score *GS*_i_ based on the weights of its associated HPO terms. The score is a sum over the ‘n’ HPO terms associated with a gene, and the weight for each term in the sum is that assigned to that HPO term in the clinical profile analysis described above.

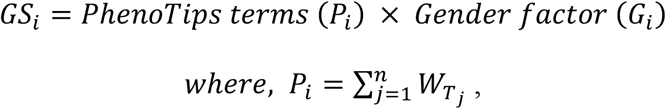

*And, G_i_ =1, if gender of phenotypic profile and genotypic profile are the same, G_i_ =0, if gender of phenotypic profile and genotypic profile are NOT the same.*

For each clinical profile, we ranked the genomes according to the highest *GS*_i_ score of any gene. The five top ranked genomes for each clinical profile were used for further analysis. If there are multiple genomes with the same score, more than five genomes will be considered for a particular clinical profile. For each of these top ranked genomes, we selected the five top scoring genes (i.e. a total of at least 25 genes per profile). There may be multiple genes with similar scores in a genome, in which case more than five may be selected. The selected genes were further filtered, removing those that do not exhibit the appropriate inheritance pattern or the appropriate ethnicity. The set of categorized variants in the remaining genes formed the set of candidate causal variants for a patient.

### Prioritized causative variants for a genome

Final prioritized causative variants were selected from the candidate set by careful manual comparison of the OMIM disease description (Hamosh et al., 2005) for the corresponding gene with each clinical profile. Variants in lower frequency bins were prioritized over those with higher frequency. For example, a novel variant in a gene will be preferred over a variant in the 0.01% frequency bin in the same gene. Confidence levels of the categories were C1 > C2 > C3 > C4 > C5 > C6. For example, a variant in a gene in Category 2 is preferred over a variant in Category 6 in the same gene. If the same gene is matched from two different genomes for a particular clinical profile, then we applied frequency and confidence criteria to select one of the two genomes.

### Probable regulatory effects of prioritized variants

In order to check for any probable regulatory effects of the prioritized variants, we noted the RegulomeDB (Boyle et al., 2012) scores which are less than 4 and so possibly part of a regulatory motif. These scores were not used for the initial prioritization of variants. A RegulomeDB score of 1a to 1f implies an eQTL. As all of the variants of interest are rare, none was found in this category. A score of 2a to 2c implies that variants at that position may directly impact a transcription factor binding site with sub-categories (2a, 2b and 2c) for different types of evidence. A score of 3a and 3b implies less strong evidence for impact on a transcription factor binding site with and subcategories (3a and 3b) indicating different types of evidence.

### Searching for Predictive Secondary variants

Here we followed the rules in ACMG (2017) (Kalia et al., 2017) to extract predictive secondary variants from 59 genes. We searched only for clinically known pathogenic and loss of function variants in those genes, as defined in Table 1 of (Kalia et al., 2017).

## RESULTS

### Demographics, clinical symptoms, and relevant genes

The SickKids5 challenge data consists of the whole genome sequencing data and clinical profiles for 24 pediatric patients, of whom 11 are male and 13 female. The age range was from five to 19 years with an average age of 10.7 years. The challenge description included the information that there are six eye disorder cases, seven neurological disorder cases, and 11 Ehlers-Danlos syndrome connective tissue disorder cases.

Notable points are that some specific HPO terms co-occur in multiple patients, some terms occur in all three classes of disease, and complex diseases co-occur with rare disease symptoms (Figure 2). Some examples: Connective tissue disorder patients exhibit symptoms involving a large number of organs such as the gastrointestinal tract including irritable bowel syndrome and Crohn’s disease (four cases), cardiovascular/hypertension (four cases), eye defects (four cases), developmental/motor delay (five cases), scarring of tissue (three cases), and bruising susceptibility (four cases). Similarly, neurological disorder patients often exhibit developmental delay or motor delay. Autism is manifested in one patient out of the seven neurological disorder cases. One neurological disorder patient is affected with an eye disorder as well as musculoskeletal disorders, including scoliosis and osteopenia. Similarly, an eye disorder patient is also affected with other musculoskeletal disorders, including hyper-extensibility of the joints and ear defects. Altogether, 10 of the 24 cases have symptoms in two or more disease classes. In total, there are 213 unique HPO terms for the 24 cases. These terms were used to compile a total 6239 potentially relevant genes from the HPO (Köhler et al., 2014) and dbNSFP (Liu et al., 2016) databases and the 319 genes in the RetNet eye disorder database (Daiger, 2004). The number of genes related to each clinical profile ranges from 350 to 4000, with an average of 1600. Supp. Figure S1 shows the number of genes for each case, grouped by disease class. Eye disorder patients have an average of 770 candidate genes. Neurological clinical profiles are usually associated with more genes, with an average of 2300 genes. Connective tissue disorder patients have the widest range, from 400 to 2800 genes.

**Figure 2:**
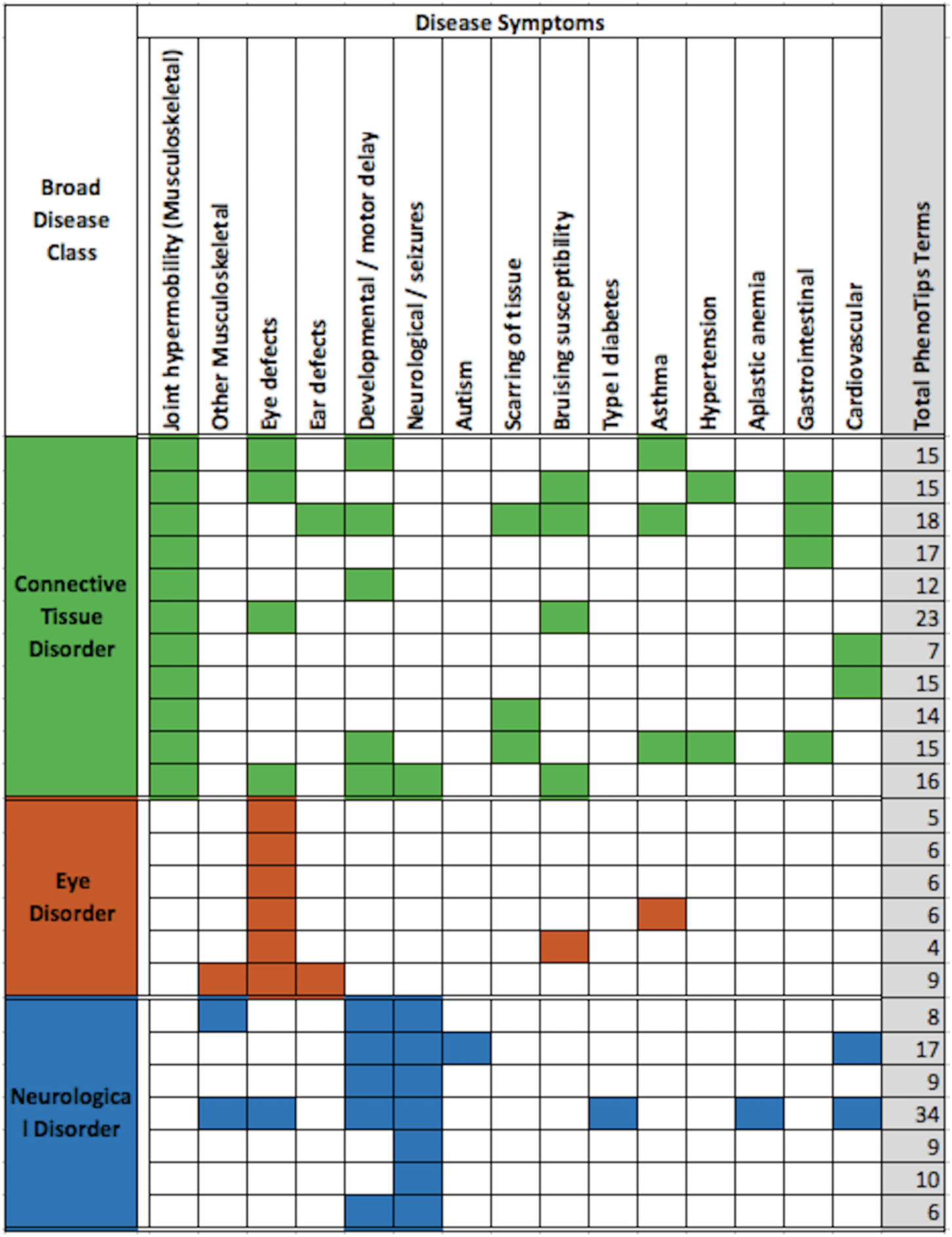
Disease classes and selected PhenoTips terms for the 24 cases. Each row shows the data for one patient and the total number of PhenoTips terms for each patient is given in the last column, shaded grey. Most patients have multiple symptoms and some symptoms occur in multiple patients. Further, some symptoms occur in all three classes of disease. Complex disease symptoms are also noted.

### SickKids5 data quality

Figure 3 shows the SickKids5 challenge data quality in terms of Ts/Tv ratio, Het/Hom alternate allele ratio, total SNV counts and rare (less than 1% population frequency) SNV counts. We compared these data with that for the corresponding ethnicities in the 1000 genome set (Auton et al., 2015) and the high quality reference Genome in a bottle (GIAB) data (Zook et al., 2016).

**Figure 3:**
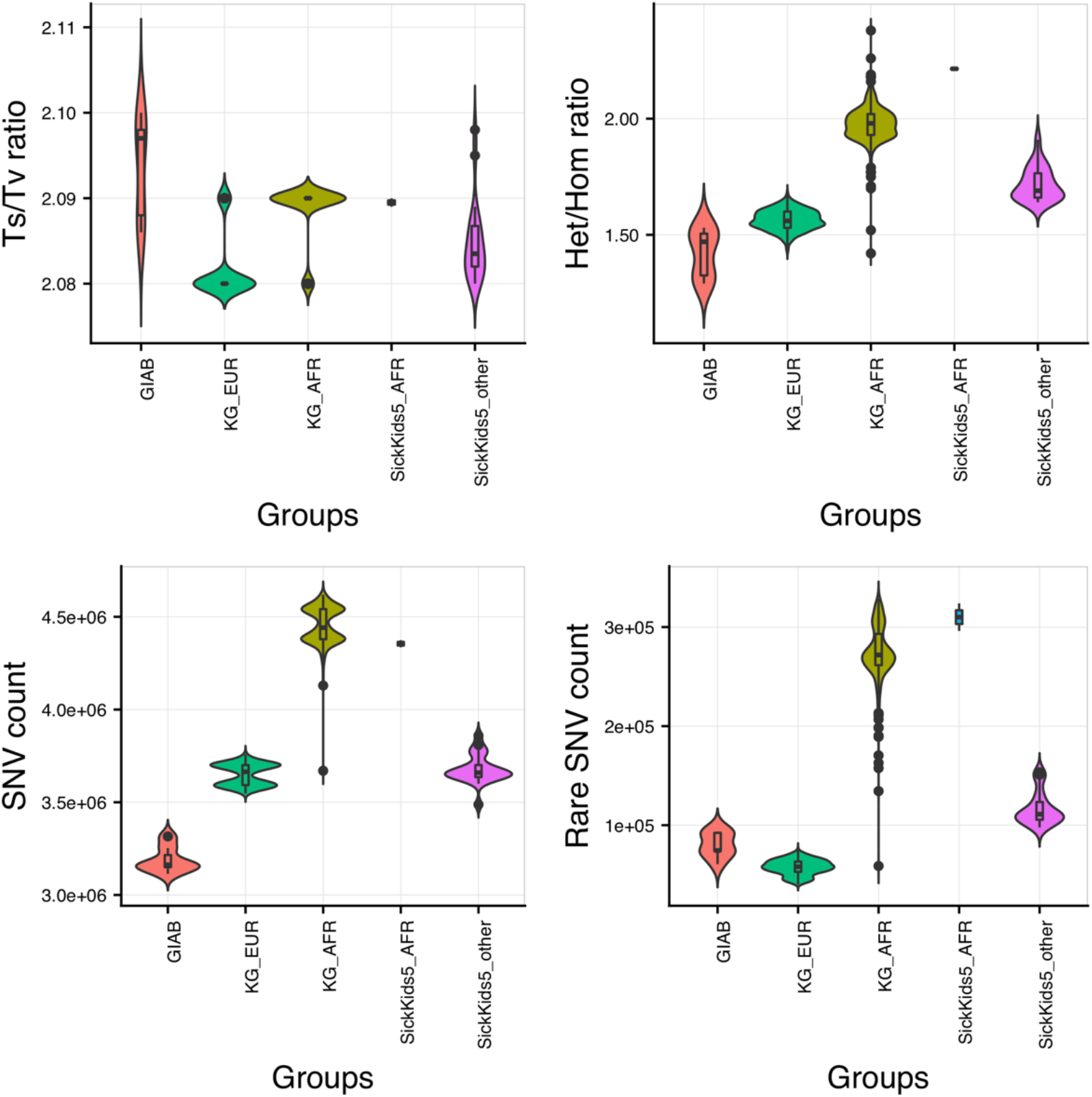
SickKids5 data quality analysis in terms of Ts/Tv ratio, Het/Hom alternate allele ratio, total SNV count and Rare (less than 1% population frequency) SNV counts in whole genomes. Only variants with ‘Pass’ status are included. 1000 genome EUR and AFR data and GIAB data provide controls. Abbreviations used: GIAB - Genome in a bottle data, KG_EUR - 1000 genome Caucasian (EUR) data, KG_AFR - 1000 genome African (AFR) data, SickKids5_AFR - SickKids5 African (AFR) data and SickKids5_other - all other SickKids5 data excluding Africans. Although there are some differences to the GIAB data controls, generally the Sickkids data appear to be of high quality.

In the previous SickKids challenge (https://genomeinterpretation.org/content/4-SickKids_clinical_genomes), we observed an excess of rare and novel variants for 25 patients with sequencing data provided by Complete Genomics (Pal et al., 2017), relative to 1000 genome data. Compared to the Complete Genomics data, the CAGI5 Illumina HiSeq X data is of better quality - the data have comparable Ts/Tv ratio, Het/Hom alternate allele ratio, and total SNV counts to that of 1000 genome data (Figure 3). Rare SNV counts in SickKids5 AFR data is comparable to that with 1000 genome AFR data. Non-AFR rare SNV counts in SickKids5 is closer to that in GIAB than 1000 genome EUR data. The excess of rare variants in both GIAB and SickKids5 data compared to 1000 genome data may be due to the increasing identification of rare variants in recent years as a result of improved sequencing technologies. Nevertheless, there is a small excess of rare as well as total variant counts for SickKids5 data compared to GIAB data, not unexpected given the very high quality of the GIAB data. To investigate this discrepancy in data quality, we checked the alternate allele fraction (alt allele counts/ref allele counts) distribution for heterozygous calls in both GIAB and SickKids5 data for all ‘PASS’ variants (Supp. Figure S2). This distribution has a broader range even within ‘PASS’ variants for SickKids5 data compared to GIAB, indicating a higher noise level in the SickKids5 data. If we restrict this alternate allele fraction distribution in SickKids5 data to the range as observed in GIAB data, the SNV counts agree.

### Distribution of candidate variants

As described in Methods, for each clinical profile, we identified the five or more top scoring (best HPO term matches) genes in the five (or more) top scoring genomes. Genes in the female 13 genomes are matched to the female profiles and genes in the 11 male genomes are matched to the male profiles. An average of 35 genes per profile were selected, resulting in a total 342 unique genes for all 24 profiles for further analysis. There is an average of five variants in each of the five genes selected in each genome, with an average total of about 116 candidate variants per clinical profile. For eye disorder clinical profiles, we also included candidate variants in the 319 RetNet genes. For each profile, the set of candidate variants were ranked using two criteria - the impact category for a variant and its frequency bin (lower frequencies rank higher) (details in ‘Categorization of variants according to their likely pathogenic impact’ section under Methods).

Figure 4 shows the counts of candidate variants in each category from selected genomes, for each profile. For all clinical profiles, the fraction of candidate variants in Category 6 (non-coding variants) is the highest (on average 83%, 5 to10 variants per included genome), followed by the variants in C2 (LOF and other high impact coding variants including missense, on average 7%, 0 to 1 variants per genome) and then in C3 category (possibly high impact missense variants, on average 5%, 0 to 1 variants per genome). Where a clinical profile contains very few HPO terms (such as eye disorder cases) with less discriminating weights amongst the terms the gene scoring scheme is less able to discriminate between genes in final reporting. This usually results in inclusion of more than five genes per genome with the same score. One such eye disorder case included an average 30 C6 candidate variants (Figure 4, last row, third column). Figure S3 shows the scores for candidate variants in the genomes selected for one clinical profile.

**Figure 4:**
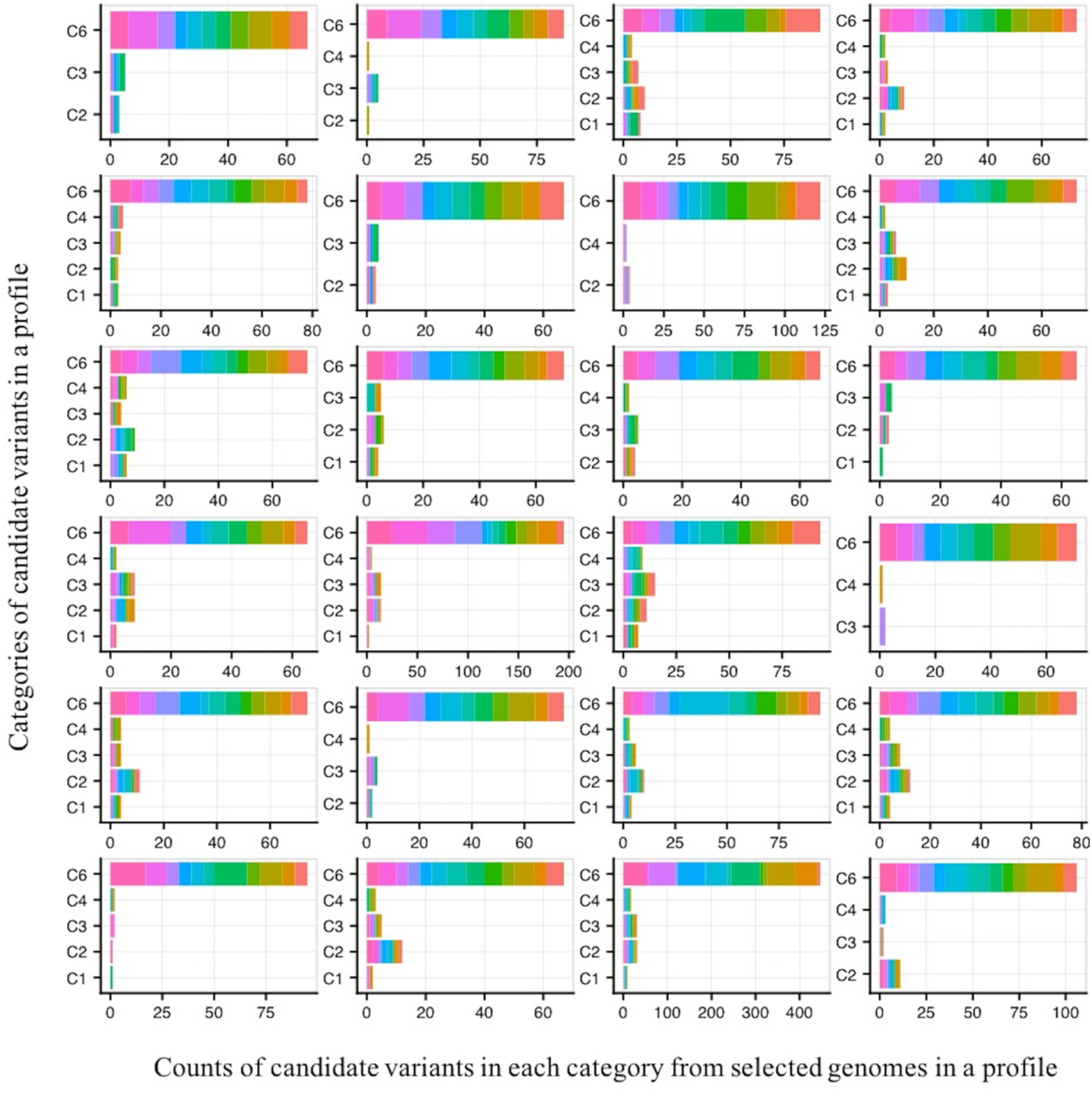
Impact distribution of selected variants for each of the 24 clinical profiles. For each profile, the number of variants falling into each of the six impact categories is shown. Variants are colored by genome of origin. The large majority of selected variants are in the C6 category: impact variants in UTRs and introns.

A subset of the candidate variants was prioritized as probably causative, based on manual inspection, as described in ‘Gene scoring scheme for selection of genomes best matching to a clinical profile’ and ‘Prioritized causative variants for a genome’ sections in Methods, including filtering by ethnicity and inheritance pattern. Comparison of the OMIM description for a gene with the clinical profile was a powerful filter, such that in most of the cases, the top scoring variant was not chosen because of a poor match. A total of 35 probable causative variants were prioritized (Supp. Table S1) for all 24 clinical profiles. Figure 5 shows the distribution of these 35 prioritized variants by category and frequency bins. 46% (16 out of 35) of the prioritized variants are in Category 6 and 44% (7 out of 16) of these are novel - that is not seen in the 1000genome, ExAc or gnomAD databases. The next highest relative occurrence of novel variants (3) and total prioritized variants (8) is in the C2 category, which includes loss of function variants together with predicted high impact non-synonymous variants. There were four prioritized variants each in the C1 and C3 categories and three variants in the C4 category.

**Figure 5:**
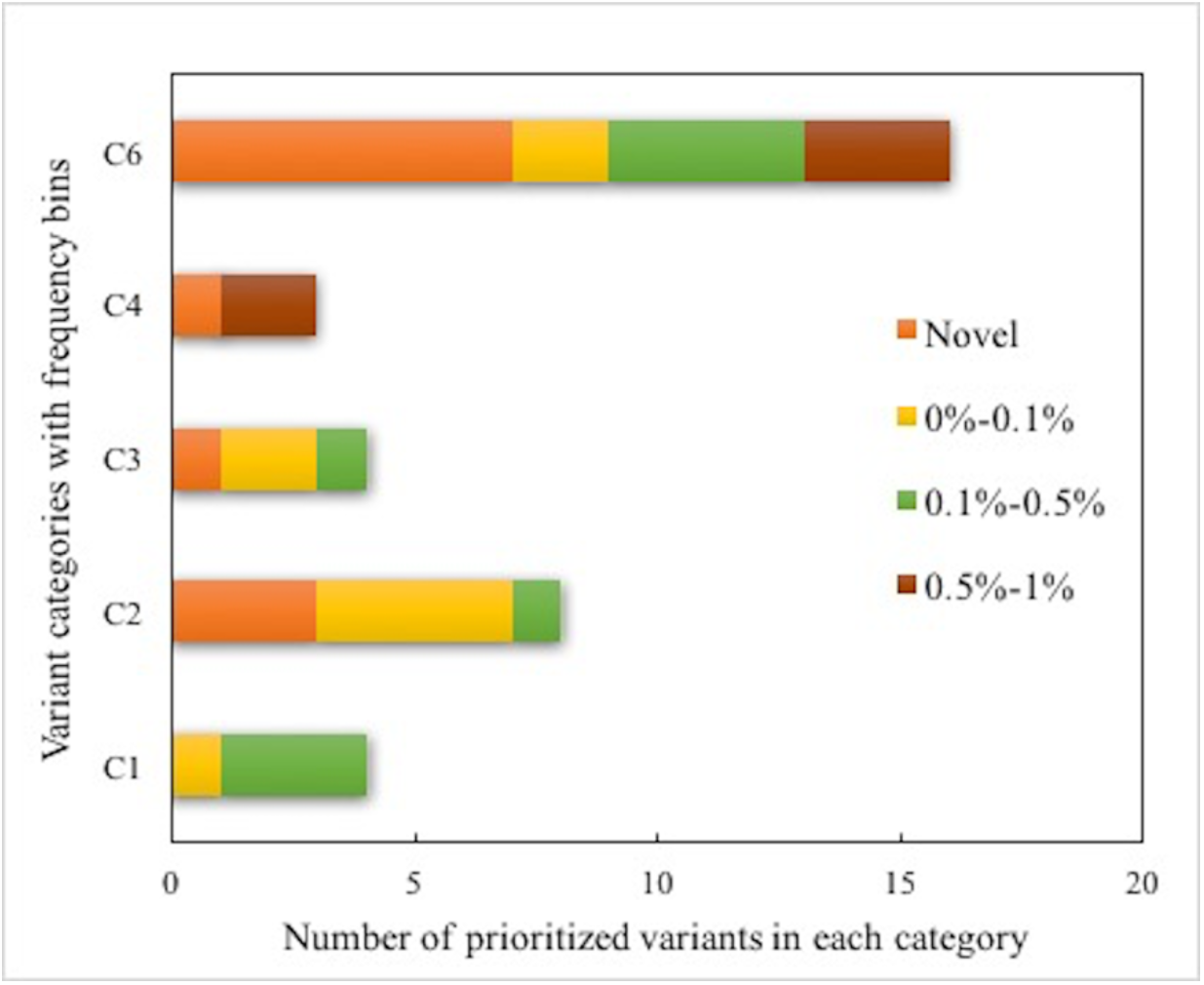
Stacked bar plot of the impact categories and frequency ranges of the 35 prioritized probable causative variants. Almost half are in the Category 6 of UTR and intronic variants.

### Molecular mechanism underlying the prioritized variants

Figure 6 shows the distribution of the 35 prioritized causative variants according to the probable underlying molecular mechanisms. 46% are missense (including those occurring in categories 1, 2, 3, and 4) and 46% are intronic or UTR variants. There are three frameshift insertion/deletion variants (9%). The missense variants have a range of impact confidence, from very high in C1 (based on clinical observation), high in C2, uncertain in C3, to predicted benign in C4. All intronic and UTR variants are predicted high impact by the Gerp++ criterion, implying conserved features at that position. Two intronic variants and one UTR variant are also predicted high impact by CADD.

**Figure 6:**
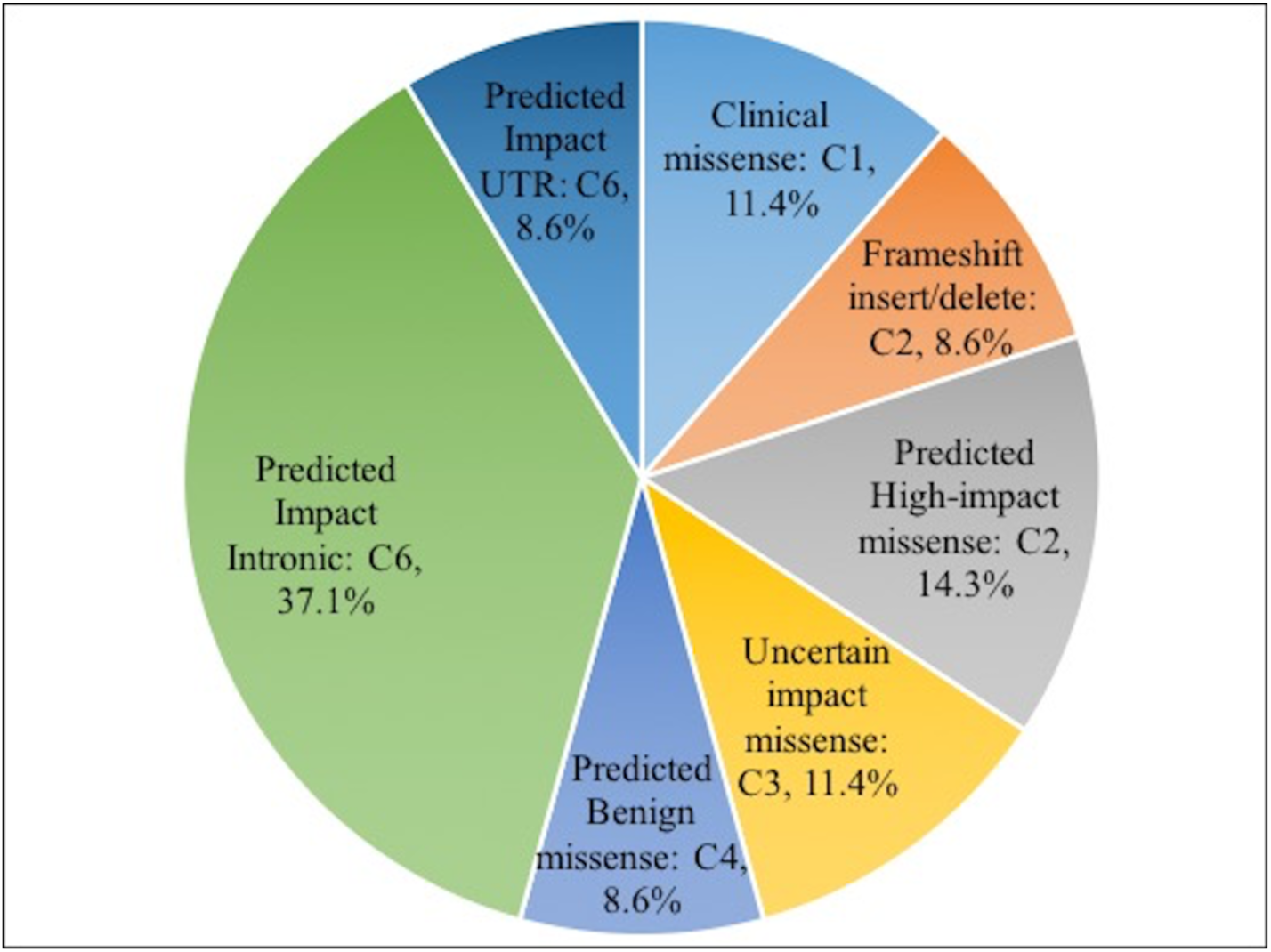
Distribution of prioritized variants by probable molecular mechanism.

Figure 7A and Figure 7B shows how the distribution of variant impact categories changed from the initial set to the candidate causative variants set to the final prioritized set. There is a 99% reduction in the number of variants going from the initial set (outermost circle in Figure 7A, all rare variants present in the genomes) to the candidate causative variants set (middle circle in Figure 7A, candidate causative variants in the selected genes). The reduction is in all three variant sets (exonic, intronic and UTR) (Figure 7B), and is a result of only a small fraction of these variants meeting the impact selection criteria from the selected genes. Exonic variants are reduced by about 92%. The lowest range of decrease is for missense variants (from 4.34 to 3.50, reduced by 85%) and loss of function variants, for example, non-frameshift indel (from 3.32 to 2.39 on the Log10 scale), frameshift indel (from 3.21 to 2.04), stop gain/loss (2.75 to 1.72). In the prioritized variants set submitted for the challenge, only missense, indels, intronic and UTR variants were selected.

**Figure 7:**
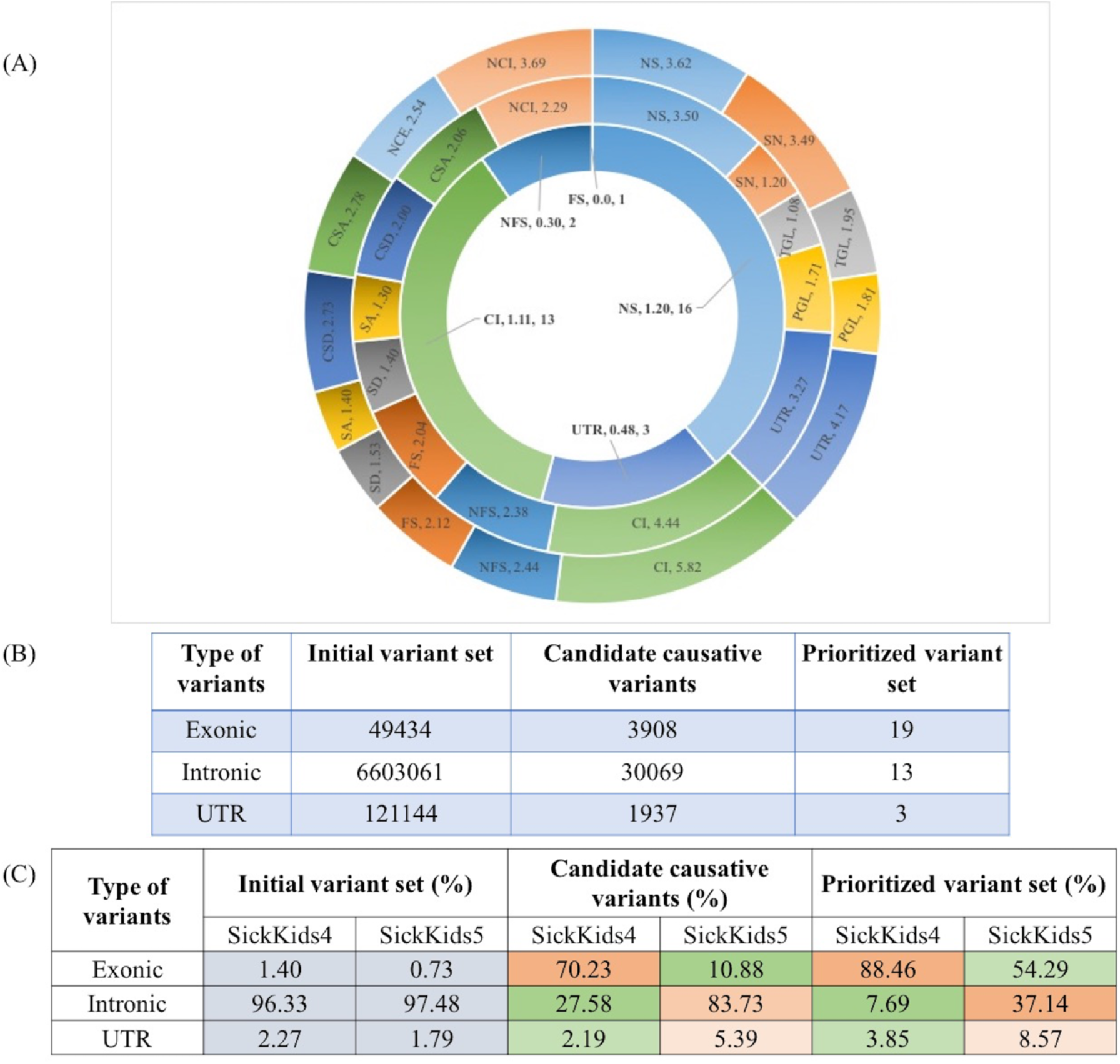
(A) Post analysis of the distribution of variant types (Log10 scale) in the total set of 6239 genes selected for the 24 SickKids5 patients. The outer most circle shows the distribution of all rare (allele frequency < 1%) variants present in the genomes. The middle circle shows the distribution of candidate causative variants in the selected genes for each clinical profile for its matching genome. The inner most circle shows the distribution of final prioritized causative variants, submitted for the challenge. Abbreviations: NS: Missense, SN: Synonymous, TGL: Start Gain/Loss, PGL: Stop Gain/Loss, UTR: UTR variants, CI: Coding Intronic, NFS: Non-frameshift indel, FS: Frameshift indel, SD: Splice donor, SA: Splice Acceptor, CSD: Close to Splice donor, CSA: Close to Splice acceptor, NCE: Non-coding exonic, NCI: Non-coding intronic. (B) The upper table shows the changes in variant composition at different stages of the selection process. (C) The lower table shows the comparison of variant composition (in %) between the SickKids4 and SickKids5 data at different stages of the selection process. The heat map highlights the differences in composition between the datasets.

We also compared the variant compositions of the SickKids5 and SickKids4 data (Figure 7C). The initial compositions are very similar, but for the candidate and prioritized variants set, there is a dramatic shift from a large majority of exonic variants in Sickids4 to a large majority of intronic variants in Sickkids5 set. The percent of UTR variants in the intronic and UTR sets also increased in SickKids5 set, by about two to three fold. This is due to the introduction of intronic and UTR variant impact predictions in the SickKids5 analysis.

### Performance in CAGI5 - matching of disease classes and exact matches

Table 1 shows the performance of the method in the SickKids5 challenge. Overall, we were able to identify the correct disease class of a genome for 12 cases and exactly matched clinical profiles to the correct genome for five cases. Disease class matching is the most successful for connective tissue disorders (6 cases, 55%), second highest for eye disorders (3 cases, 50%) and least successful (3 cases, 43%) for neurological disorders. The five correct profile/genome matches are composed of two connective tissue disorder cases, two for eye disorders, and one for neurological disorder.

According to the data provider (who was also the challenge assessor), out of the five exact match cases, the genes carrying two eye disorder diagnostic variants and one of the genes for connective tissue disorder diagnostic variants are possibly correct (Table 1 and Supp. Table S1). The eye disorder diagnostic variants are: (1) Compound heterozygous coding-intronic variants (conserved by Gerp++ scores) in the *USH2A* gene - annotated for recessive retinitis pigmentosa and (2) compound heterozygous variants, one clinical missense variant and a coding-intronic variant (conserved by Gerp++ score) in the *ABCA4* gene - annotated for retinitis pigmentosa, rod-cone dystrophy and other eye disorders. For the connective tissue disorder case we prioritized two heterozygous variants in the *FBN1* gene, for an autosomal dominant inheritance pattern (information not provided in the clinical profile). One variant is in the 3’ UTR region, conserved by Gerp++ score, and the other is a novel coding-intronic variant, conserved by Gerp++ and an impact variant according to CADD. We prioritized both variants as either might be the correct diagnostic variant, and we could not distinguish between them. The non-coding variants were checked for possible regulatory effects with RegulomeDB. The *FBN1* intronic variant has a score of 3a, indicating partial evidence for transcription factor binding (the RegulomeDB annotation was not used in the variant selection procedure). Variants in the other two genes (*ABCA4* gene and *USH2A* gene) have RegulomeDB scores greater than 4 (implying lack of evidence for the variant disrupting the transcription factor binding site).

### Illustrative example of matching a genome to a phenotypic profile

Clinical profile N is of an 11-year-old female whose indication for referral is ‘Cerebral arteriovenous malformation’. The ‘Clinical symptoms and physical findings’ section for this patient also note ‘aortic dilation’ and ‘joint hypermobility’, both described as ‘borderline’. In the subjective weighting of these HPO terms, we put the highest weight on ‘Cerebral arteriovenous malformation’, a lower weight on ‘joint hypermobility’ and related HPO terms and a further lower weight on ‘aortic dilation’ related terms. The least weight was set for the neurological ‘Headache’ symptom. With these weights, we selected the top scoring genomes for profile N. In this case, there were many equally scoring genes, resulting in all 13 female genomes being selected. Three of these genomes contained the same three highest scoring genes: *ACVRL1, ENG,* and *SMAD4* with matching terms for only ‘Headache’ and ‘Cerebral arteriovenous malformation’. The next highest scoring gene was *FBN1* with matching terms for ‘joint hypermobility’ and ‘aortic dilation’. *FBN1* was considered more relevant (according to the OMIM disease description for Marfan syndrome which matches the presumed disease class of connective tissue disorder in the profile) than any of the three top scoring genes. So we selected all variants in that gene for further analysis. We found *FBN1* variants in a total of seven genomes. These are all category 6 variants, falling in the UTR and intronic regions. The frequency criteria were used to select the final variant. The variants span all four frequency bins (described in ‘Categorization of variants according to their likely pathogenic impact’ section under Methods). Two are novel, and so were given priority. One of these, in genome 056, is annotated as pathogenic by two methods (GERP++ and CADD) and on that basis we selected genome 056 for clinical profile N. This is a case where we successfully matched the clinical profile with the correct genome.

### Puzzling cases - limitation of phenotype-weighted scoring

One of the most critical factors in the phenotype-weighted scoring strategy is to correctly rank the importance of symptoms in a clinical profile, otherwise the predictions will be erroneous. This information, which is usually obvious to physicians, is typically absent from the clinical profile documentation. In the SickKids5 challenge, we considered the ‘indication for referral’ field to understand the relative importance of clinical symptoms. We failed to identify the proper disease class for one neurological disorder patient (J) because the indication for referral was ‘mitochondrial disorder’ and this patient also has multi-organ failure, including severe eye problems, seizures, and connective tissue disorders. For this patient, we found a ClinVar variant in chromosome M, with a disease description very similar to that of the patient. Supplementary Table S2 documents this and two other puzzling cases where high confidence loss of function variants are not causative. In one, for clinical profile ‘I’, we found a non-frameshift-deletion variant in the *ELN* gene for one genome and in the same gene we found a 5’ UTR variant in a different genome. According to our prioritization criteria, we selected the loss of function variant (non-frameshift-deletion) as the causative variant. However, the genome with the 5’ UTR variant in *ELN* was the correct match.

### Re-evaluation of the genome to clinical profile matches in the post-submission phase

Availability of the answer key in the post-submission phase allowed us to examine the genes in the correct genome more critically for the matched profile, followed by prioritizing suitable impact variant(s) as we did for the challenge. Supp. Table S3 lists 44 such prioritized variants in 26 genes for 24 SickKids5 patients. As these are all cases where the conventional bioinformatics pipeline did not identify diagnostic variants (implying no suitable clinical variant or loss of function variants or coding variants of unknown significance were found), we expected the task to be difficult. Often it seems that there are disparate symptoms that can only be accommodated by potential causative variants in two different genes, rather than one. One such example is for clinical profile F, where for connective tissue disorder, we identified a rare coding-intronic variant in the *COL5A2* gene. However, this patient also has very fragile skin and a food intolerance problem. We identified another novel 5’ UTR5 variant in the *PLEC* gene consistent with these additional symptoms.

Supp. Figure S4 shows that the fraction of non-coding variants (intronic and UTR) is much higher in the post-submission analysis (46% in submitted predictions vs. 77% in post-submission predictions) with 38% (13 out of 34 C6 category variants) being novel. Accordingly, the missense variant fraction is reduced to 20% compared to 46% in the submitted predictions. Validation of these non-coding variants is difficult. In order to check for any probable regulatory effects of these variants, we noted the RegulomeDB scores less than 4 (see the ‘Probable regulatory effects of prioritized variants’ section under Methods) in Supp. Table S3. Out of 44 prioritized variants, 36 returned a RegulomeDB score less than 4. We found three variants with a score of 2 and two variants with a score of 3 out of the 36 variants. These variants are: A neurological disorder case with a score of 2a (rare - less than 0.05% allele frequency, intronic variant in *CHD2*, related to myoclonic encephalopathy). Two connective tissue disorder cases with score 2b - one is the novel 5’ UTR variant in the *PLEC* gene (described above) and another one is a rare - less than 0.01% allele frequency - missense variant in the *TNXB* gene, predicted to be deleterious by half of the methods used (so a C3 category variant). We found one neurological variant with score 3a (novel coding-intronic variant in *ARID1B*, related to developmental delay with seizures). Another novel connective tissue disorder variant in the *FBN1* gene with score 3a was already included at the challenge stage (described in the ‘Performance in CAGI5’ section).

### Predictive secondary variants

Supplementary Table S4 lists the eight predicted secondary variants we submitted, found in six SickKids5 patients. There are three novel secondary variants (a clinical missense variant in *KCNH2* for Long QT syndrome, a non-frameshift-deletion in *MSH2* for Lynch syndrome, and a non-frameshift-deletion in *BRCA2* for hereditary breast cancer) found in one neurological disorder patient (of African origin). The other predicted secondary variants are clinical variants in the *MSH6* and *MSH2* genes for Lynch syndrome and in the *LMNA* gene for hypertrophic cardiomyopathy. The same novel *MSH2* variant was found in two patients (genome 081 and 091) and according to the challenge assessor (Kasak et al., 2019), these might be sequencing errors. The alternate allele fraction (alt allele counts/ref allele counts) of these variants (Supp. Table S4) are poor, 0.36 and 0.42 for genomes 081 and 091 respectively, supporting the sequencing error hypothesis.

## DISCUSSION

The CAGI SickKids5 challenge provided an opportunity to assess methods for correlating whole genome sequencing data to clinical information. Participants were asked to predict the disease class (eye, neurological and connective-tissue disorders) of 24 undiagnosed whole genomes and to identify which genome matches to each clinical profile. To address this challenge, we developed a semi-automated gene-centric method. The method builds on one we had previously implemented for identifying causative variants based on clinical information in the CAGI4 SickKids challenge (Pal et al., 2017). The key CAGI5 innovation is the introduction of a phenotype weighting scheme to evaluate the match of gene descriptions and clinical profiles, using HPO (Human Phenotype Ontology) (Köhler et al., 2014) terms. Using this approach, we were able to identify correct disease classes for 12 of the 24 genomes and to match five genomes to the correct clinical profiles. Analysis of the method’s performance and results have provided a number of insights into issues related to the scoring scheme, nature of prioritized variants, methodology used, and key factors in extracting clinical information from a whole genome.

### Phenotype-weighted scoring scheme for genes

Success of the phenotype-weighted scoring scheme depends on how effectively the clinical documentation portrays patients’ symptoms. SickKids clinical profiles are constrained to terms in the Human Phenotype Ontology, and phenotypes associated with specific genes are also available in that form. Thus, HPO based gene by gene matching to a clinical profile provides a strategy for the selection of genes that are most likely to harbor causative variants. However, simply looking for overlap between gene and profile HPO terms is a not sensitive enough matching algorithm. Instead, we assigned a weight to each of the clinical profile terms, depending on the prominence of the term in the description (for example, up-weighting referral terms), and down-weighting terms that are characterized as less severe and those that do not match the disease class.

Although this approach did allow us to match a significant number of genomes to profiles, it has issues in some specific circumstances. Generally, too few terms in a clinical profile are not informative enough. For example, in one eye disorder case (case W), there were only two effective HPO terms in the profile, resulting in low discriminatory power and the selection of a large number of genes. As a consequence, there are a very large number of candidate variants (Figure 4, last row and third column). Although more terms are usually better, term combinations are of varying discriminatory power. For example, in one neurological disorder (case J), the patient has HPO terms for all three disease classes. As a result, we failed to identify the correct disease class and so did not assign appropriate weights, resulting in an erroneous choice of genome for the profile. A limitation of the current scoring method is that it does not penalize for missing terms - ones that are not present for a gene but are in a profile or conversely ones that are not present in a profile but are there for a gene. For example, there are some genes related to eye disorders that are also related to hearing problems, and some that are not. The method as used in CAGI5 would select all these genes even if the profile includes hearing HPO terms (for example, case X). These limitations will be addressed in future versions of the method.

### Nature of prioritized variants

In SickKids4 (Pal et al., 2017) we mostly prioritized coding variants (88% of all types of variant in SickKids4 vs. 54% in SickKids5, Figure 7C). The high proportion of non-coding candidate variants in SickKids5 is a consequence of introducing two more non-coding variant impact analysis methods, GERP++ (Davydov et al., 2010) and Eigen (Ionita-Laza et al., 2016), in addition to CADD (Kircher et al., 2014) which was also used in CAGI4. GERP++ turned out to select many more variants than CADD, whereas Eigen returned none. While CADD for coding missense variants is considered to have a reasonable performance (Anderson & Lassmann, 2018), CADD scores for non-coding variants have been found to have limited clinical utility in one study of rare non-coding variants in a hereditary cancer panel (Mather et al., 2016). There is no such benchmarking data for rare non-coding variants available for GERP++ scores. The authors of the method report an overall very low (0.86% in (Davydov et al., 2010)) false positive rate. A general problem at present is that methods for non-coding variants are less mature than those for coding. Nevertheless, non-coding variants do play a critical role in our analysis (four out of five exact match cases in our predictions were identified using non-coding variants).

### Scope for improvement

The cases in the Sickkids5 challenge were unresolved by a traditional pipeline. Although we and another group did better than random at matching genomes to profiles and the assessors considered some prioritized genes promising, most of the cases remain a mystery. And, as noted earlier, in general, rare disease pipelines have a success rates below 50% (Clark et al., 2018). There are number of possible explanations for the low yield of diagnostic variants, even given whole genome sequencing data. We conclude by considering the most relevant of these, and the prospects for progress:

(A) New genes related to specific disease phenotypes are continually being discovered (Friedman et al., 2019; Guelfi et al., 2019) implying that there are many more still to be found. A strategy that might help address this problem is to consider all putative impact variants in all genes, and see if any of these genes have phenotype descriptions that offer some clue to a possible match (a genotype to phenotype approach (Hu et al., 2013; Wang et al., 2010)). As more rare disease genome data accumulate, it will be possible to look for enrichment of impact variants in particular genes in the presence of particular phenotypes, and this is likely to prove a powerful approach providing a long-term solution. In the meantime, for analysis of a single genome, and even more so for the Sickkids5 challenge with 24 genomes, the large number of putative impact variants makes the strategy very difficult. If we consider only C2 variants - not clinically recognized but confidently predicted high impact, there are an average of about one in every 8 genes, so that in a single full genome there will be about 3000 variants to screen. When considering 24 genomes, there will be about 70,000 such variants. Nevertheless, it may be possible to develop a tuned version of the phenotype scoring scheme we used in the challenge to filter the variants. Consideration of knockout or knock in data in model organisms (Smedley et al., 2015) together with such clues may be partially effective.

(B) In some complicated clinical profiles (such as for connective-tissue disorders or neurological disorders), contributions from more than one gene may be present. Indeed, one study estimates that this occurs in 5% of rare disease cases (Y. Yang et al., 2014) and this likely is a considerable underestimate. We see evidence for involvement of multiple genes in three cases (Supp. Table S3). For example, for clinical profile F, a connective tissue disorder, we originally predicted a novel missense variant in *EP300* as causative, with an OMIM disease description of Rubinstein-Taybi syndrome. According to the assessor, even though we selected the correct genome, this disease description is not an adequate match to the patient’s symptoms. On further inspection of the genome, knowing it is a correct match, we found an intronic variant in *COL5A2*, which is related to the classic Ehlers-Danlos syndrome (a partial match of patient’s symptoms). We also found two other novel impact UTR variants in *PLEC* (the gene has an autosomal recessive inheritance pattern in OMIM), with an OMIM disease description of epidermolysis bullosa with pyloric atresia, related to the patient’s fragile skin and food intolerance symptoms.

(C) As discussed above, present methods for identifying non-coding impact variants are not mature. Recent strong CAGI results for predicting which variants affect expression are encouraging in this regard (Shigaki et al., 2019), and it would be interesting to see how some of the more successful methods used there perform on the SickKids data.

(D) Non-standard descriptions in clinical reports: An advantage of the Sickkids data is the use of HPO terms to describe patients’ symptoms (Girdea et al., 2013). That greatly facilitated identification of candidate genes, and its broader adoption by other analysis centers will improve performance. In addition, some kind of weighting scale would also help - it may be obvious to a physician that a particular HPO is not central to a patient’s phenotype, but at present that information is often not available in the record.

(E) The role of variants affecting so far poorly understood function, particularly those that may affect chromatin structure. Examples of these have already been found in cancer (Fudenberg & Pollard, 2019; Makova & Hardison, 2015). It is not clear how well general non-coding impact methods will work on such variants, and they may be very far from genes, requiring a much larger total number of variants to be considered, with an accompanying rise in false positives. Advances in resolving three-dimensional chromatin structure and how it varies (Kishi & Gotoh, 2018; Marti-Renom et al., 2018; Qi & Zhang, 2019) hold long term hope for progress here.

## ACKNOWLEDGEMENTS

We are grateful to Dr. Stephen Meyn at the Hospital for Sick Children, Toronto for generously making the challenge data set available.

## Data availability statement

The prediction files are available to registered users from the CAGI web site (https://genomeinterpretation.org/SickKids5_clinical_genomes). Restrictions apply to the availability of the genomic data, which were used for this study under license from the Hospital for Sick Children. Genomic data are available from Dr. Stephen Meyn with the permission of the Hospital for Sick Children.

