## Supplementary material for "Matching whole genomes to rare genetic disorders: Identification of potential causative variants using phenotype-weighted knowledge in the CAGI SickKids5 clinical genomes challenge": Supp. Fig S1, Supp. Fig. S2, Supp. Fig. S3, Supp. Fig. S4, Supp. Table S1, Supp. Table S2, Supp. Table S3, Supp. Table S4

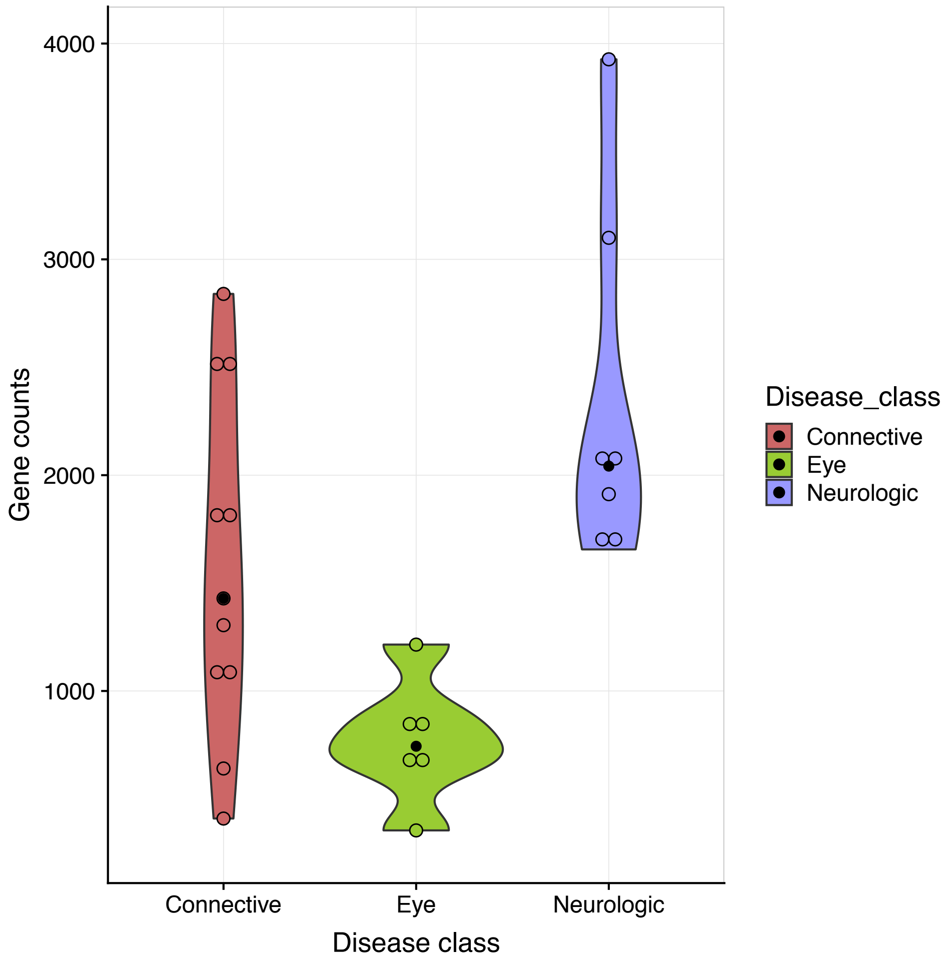

Figure S1: Violin plot for gene counts extracted for each phenotypic profile, grouped by the three broad disease classes. Open circles show individual patient gene counts. Black dots show median values.

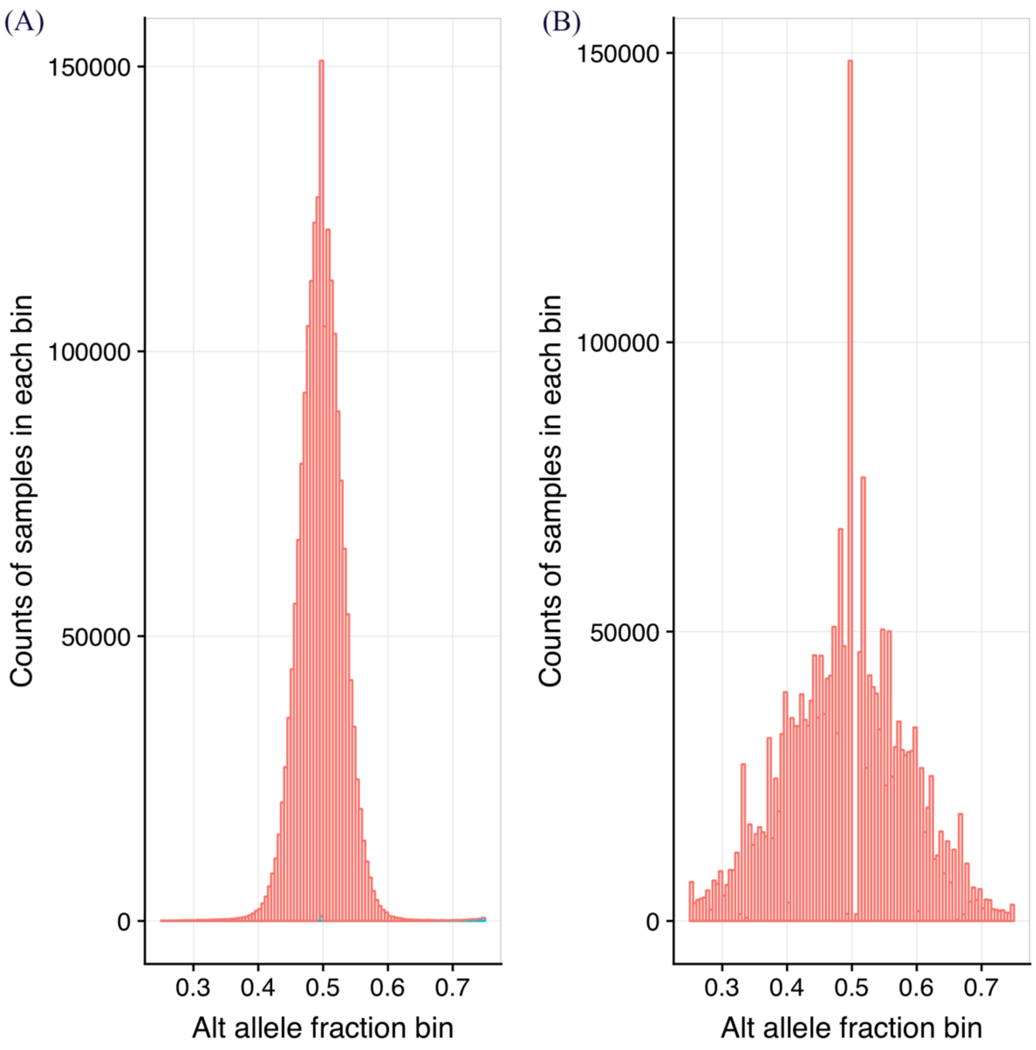

Figure S2: Alternate allele fraction distribution for (A) The Caucasian GIAB heterozygous data and (B) An example Caucasian SickKids5 patient. SickKids5 allele distributions are wider, indicating data with higher noise levels.

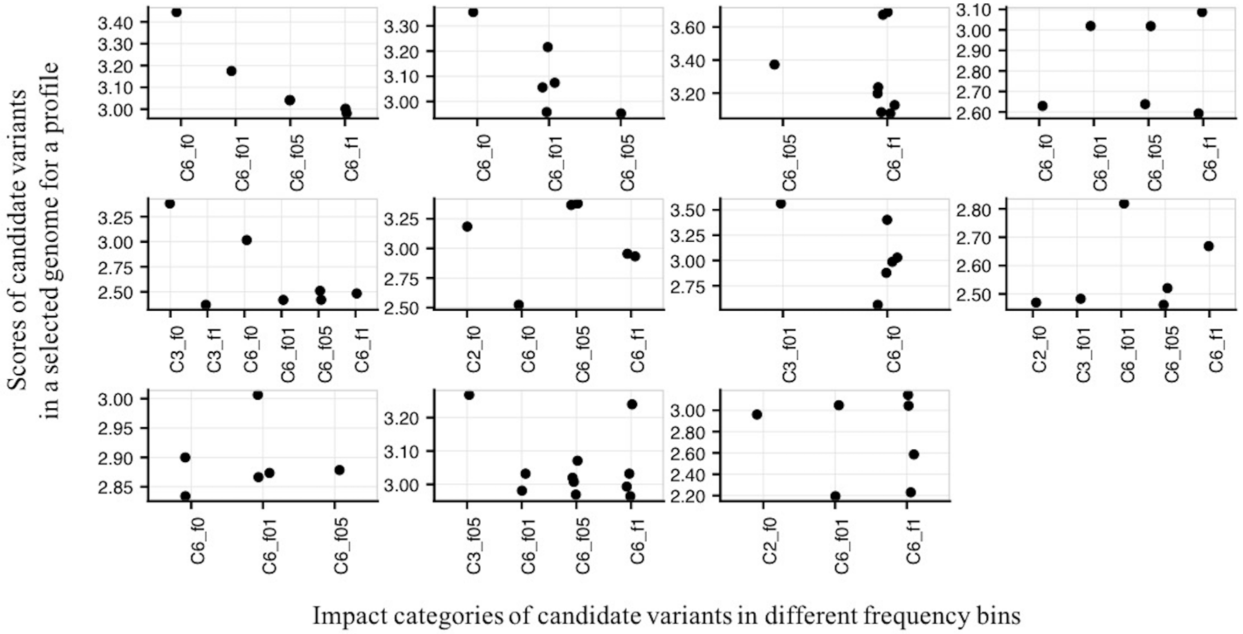

Figure S3: Candidate variant scores in selected genomes for an example clinical profile. This individual is male, and there is degeneracy in their genomic scores. As a result, variants in all 11 male genomes were considered. Each plot shows the scores of candidate variants in one of the 11 genomes. The Y axis shows the HPO match score between the gene each variant lies in and the patient's clinical profile. The X axis shows category and frequency bins for the candidate variants - the category name is followed by frequency bin information, where f0 represents a novel variant, f01 variants with population frequency greater than 0 and less than 0.01%, f05 for that greater than 0.01% and less than 0.05% and f1 is for variants greater than 0.05% and less than 1% population frequency.

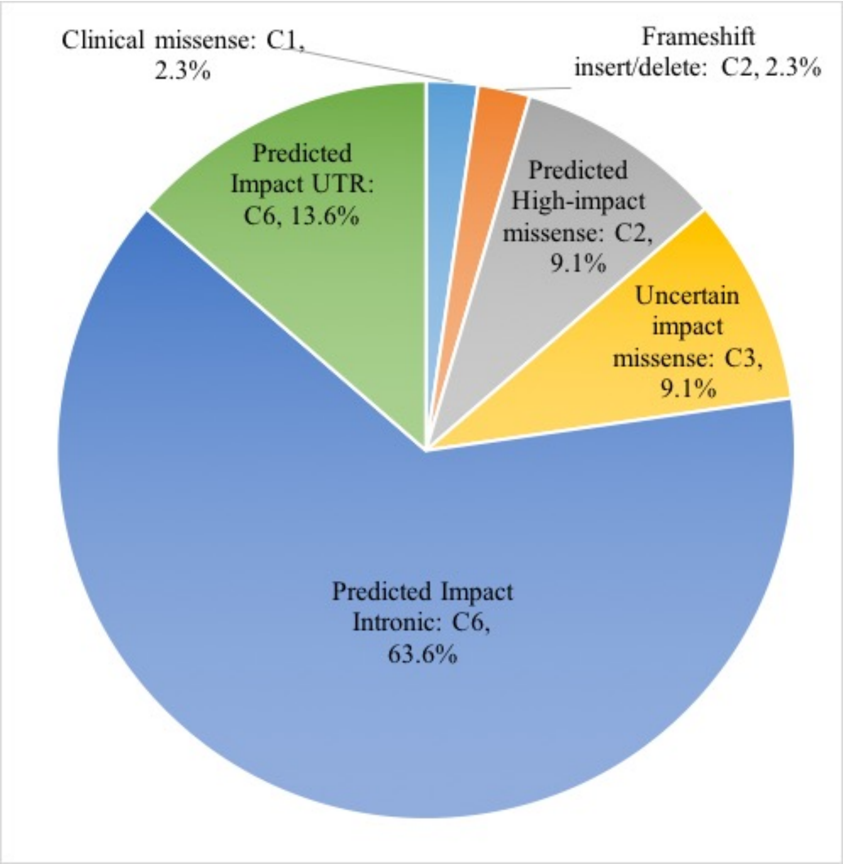

Figure S4: Post-Submission analysis - distribution of prioritized variants in different categories according to their probable molecular mechanism.

**Supplementary Table S1:** Submitted diagnostic variants for the SickKids5 challenge.

| Genome | Submitted assignment of patient | Submitted disease class | Gender | Submitted gene | Submitted probable diagnostic variants | Mechanism involved | Max minor allele frequency | OMIM diseases | Comments |
|---|---|---|---|---|---|---|---|---|---|
| **007** | G | Neurologic | F | PCLO | 7:82584646:C:T, 7:82384308:C:T | Nonsyn_5/6, C2_f01; UTR3_1/3, C6_f0 | 0.001 in 1000 genome, Novel | Pontocerebellar hypoplasia, type 3 | Compound het variants |
| **009** | O | Eye | F | TRPM1 | 15:31318422:C:G | Nonsyn_3/7, C3_f05 | 0.004 in ExAC | Nyctalopia;Abnormal electroretinogram;Retinal dystrophy | Missense variant |
| **017** | H | Eye | M | ABCA4 | 1:94568686:C:T, 1:94470320:C:T | Clinical_HGMD_Nonsyn_C1_f05, Intronic_1/3,C6_f01 | 0.002 in gnomad, 0.006 in gnomad | Retinitis pigmentosa, rod-cone dystropy and other eye disorders | Nonsyn variant is clinical variant by HGMD (DM?) and other one is impact variant by Gerp++, which is also |

| | | | | | | | | | |
|---|---|---|---|---|---|---|---|---|---|
| | | | | | | | | | shared by one more patient. But both of these variants are only present in H. |
| **018** | M | Eye | M | PROM1 | 4:16010674:A:G | Nonsyn_6/7, C2_f01 | 0.000085 in gnomAD | Retinitis pigmentosa 41;Macular dystrophy, retinal, 2 | Missense variant |
| **021** | T | Connective | F | FLNA | X:153579297:T:C | Benign_Nonsyn, C4_f0 | Novel | Heterotopia, periventricular;Cardiac valvular dysplasia, X-linked;Intestinal pseudoobstruction, neuronal;Frontometaphyseal dysplasia 1;Melnick-Needles syndrome;Terminal osseous dysplasia;Otopalatodigital | Novel missense |
| **030** | K | Connective | M | ADAMTS2 | 5:178549719:G:A,5:178614102:G:C | Benign_Nonsyn, C4_f1; | 0.007 in ESP, 0.005 | Ehlers-Danlos syndrome, | Compound het variants |

| | | | | | | | | | |
|---|---|---|---|---|---|---|---|---|---|
| | | | | | | Intronic_1/3, C6_f1 | in 1000 genome | dermatosparaxix type | |
| **039** | R | Neurologic | M | WWOX | 16:78868278:C:T,16:79056317:C:T | Intronic_1/3, C6_f05; Intronic_1/3, C6_f0 | 0.004 in 1000 genome, Novel | Epileptic encephalopathy, early infantile, 28;Spinocerebellar ataxia, autosomal recessive 12 | Intronic variants by GERP++ and CADD |
| **042** | V | Connective | F | COL1A2, TNXB | 7:94055784:C:A, 6:32032628:C:G | Nonsyn_1/7, C3_f01; Benign_Nonsyn, C4_f1 | 9.87E-05 in gnomAD, 0.009 in 1000 genome | Osteogenesis imperfecta, type II;Osteogenesis imperfecta, type IV;Osteoporosis, postmenopausal;Ehlers-Danlos syndrome, cardiac valvular type;imperfecta, type III | Two missense variants, AD mode |
| **056** | N | Connective | F | FBN1 | 15:48700642:T:C, 15:48849792:C:A | UTR3_1/3, C6_f01; Intronic_2/3, C6_f0 | 0.00003228 in gnomAD, Novel | Marfan syndrome | UTR3 variant is impact variant by GERP++. The novel intronic variant is impact variant by both Gerp++ |

| | | | | | | | | | |
|---|---|---|---|---|---|---|---|---|---|
| | | | | | | | | | and CADD. |
| **057** | E | Neuro logic | F | RAI1 | 17:17696987:C:T | clinical_hgmd_N onsyn, C1_f05 | 0.003 in ExAC | Smith-Magenis syndrome | Clinical variant |
| **067** | L | Conne ctive | M | FBN1 | 15:48737684:T:C | nonsyn_2/8, C3_f0 | Novel | Acromicric dysplasia;Marfan syndrome;Ectopia lentis, familial;Geleophy sic dysplasia 2;Marfan lipodystrophy syndrome;Stiff skin syndrome;Weill-Marchesani syndrome 2, dominant | Novel missense |
| **068** | S | Conne ctive | F | LMNA | 1:156105054:G:T | Nonsyn_7/7, C2_f0 | Novel | Restrictive dermopathy, lethal;Cardiomyo pathy, dilated, 1A;Muscular dystrophy, congenital;Lipody strophy, familial partial, type 2;Heart-hand syndrome, Slovenian type;Emery-Dreifuss muscular dystrophy | Novel missense |

| | | | | | | | | | |
|---|---|---|---|---|---|---|---|---|---|
| **071** | U | Neuro logic | M | WWOX | 16:78827367:T:C ,16:78864508:C: G | Intronic_1/3, C6_f05; Intronic_2/3, C6_f05 | 0.003 in 1000 genome, 0.004 in 1000 genome | Epileptic encephalopathy, early infantile, 28;Spinocerebella r ataxia, autosomal recessive 12 | Compoun d het intronic variants by GERP ++ and CADD |
| **076** | D | Conne ctive | F | COL5A1 | 9:137722010:G: GC | FrameShiftInsert, C2_f0 | Novel | Ehlers-Danlos syndrome, classic type, 1 | EDS variant |
| **078** | Q | Conne ctive | F | COL1A2 | 7:94049587:C:T | Nonsyn_7/7, C2_f01 | 0.000008 in gnomAD | Osteogenesis imperfecta, type II;Osteogenesis imperfecta, type IV;Osteoporosis, postmenopausal;E hlers-Danlos syndrome, cardiac valvular type;imperfecta, type III | Missense variant |
| **079** | A | Conne ctive | M | SKI, FBN1 | 1:2234817:C:T,1 5:48855195:A:G | Nonsyn_3/7, C3_f01 ; Intronic_1/3, C6_f0 | 0.000004 in gnomAD, Novel | Shprintzen-Goldberg syndrome; Acromicric dysplasia;Marfan syndrome;Ectopia lentis, familial;Geleophy sic dysplasia 2;Marfan lipodystrophy syndrome;Stiff | Missense and intronic, by GERP++ |

| | | | | | | | | | |
|---|---|---|---|---|---|---|---|---|---|
| | | | | | | | | skin syndrome;Weill-Marchesani syndrome 2, dominant | |
| **081** | X | Eye | F | BRAF | 7:140437592:G:A | Intronic_1/3, C6_f05 | 0.002 in 1000 genome | LEOPARD syndrome 3;Noonan syndrome 7;Cardiofaciocutaneous syndrome | Intronic variant by GERP++ |
| **091** | J | Neurologic/Eye | F | POLG | 15:89873364:C:G,15:89876827:TTGCTGC:T,M:7730:A:G | clinical_hgmd_Nonsyn, C1_f05; NonFrameShiftDelete, C2_f0 | 0.004 in ESP, Novel | Progressive external ophthalmoplegia, autosomal recessive 1;Mitochondrial DNA depletion syndrome 4B (MNGIE type);Progressive external ophthalmoplegia, autosomal dominant 1;Mitochondrial | Found one mitochondrial diagnostic variant related to this disease and compound het POLG diagnostic variants |
| **092** | I | Connective | F | ELN | 7:73470620:AGTTGGAGGCATTCCTACTTACGGG:A | NonFrameShiftDelete, C2_f01 | 0.001 in ESP | Supravalvar aortic stenosis;Cutis laxa, autosomal dominant | LOF variant |
| **093** | F | Connective | M | EP300 | 22:41574743:A:G | Nonsyn_4/6, C2_f0 | Novel | Rubinstein-Taybi syndrome 2 | Novel missense |

| | | | | | | | | | |
|---|---|---|---|---|---|---|---|---|---|
| **095** | C | Eye | M | USH2A | 1:215953583:A:G, 1:215964830:T:G | Intronic_1/3, C6_f05; Intronic_1/3, C6_f0 | 0.002 in 1000 genome, Novel | Retinitis Pigmentosa | Recessive trait |
| **097** | W | Eye | F | IMPDH1 | 7:128040571:G:A | Clinical_hgmd_Nonsyn, C1_f01 | 0.000097 in gnomAD | Retinitis pigmentosa 10 | Missense variant |
| **099** | B | Neurologic | M | PIGT | 20:44044789:C:T | UTR5_2/3, C6_f01 | 0.000016 in gnomAD | Paroxysmal nocturnal hemoglobinuria 2;Multiple congenital anomalies-hypotonia-seizures syndrome 3 | UTR variant |
| **102** | P | Neurologic | M | EPM2A | 6:145992717:A:C, 6:145983098:A:G | Intronic_1/3, C6_f0; Intronic_1/3, C6_f0 | Novel, Novel | Epilepsy, progressive myoclonic 2A (Lafora);Epilepsy, progressive myoclonic 2B (Lafora) | Compound het variants |

**Supplementary Table S2:** Puzzling cases in submitted predictions for the SickKids5 challenge

| **Wrong Clinical profile** | **Wrong genome** | **Disease class** | **Apparent wrong prediction of probable diagnostic variant** |
|---|---|---|---|
| **J** | 091 | Neurologic | Clinical variant and nonFrameshift delete in POLG, ClinVar mitochondrial variant |
| **D** | 076 | Connective | Frameshift Insert in COL5A1 |
| **I** | 092 | Connective | Non-frameshift delete in ELN |
| **Correct Match** | | | |
| **I** | 081 | Connective | UTR5 variant in ELN |

**Supplementary Table S3:** Post-submission analysis of diagnostic variants, with knowledge of the answer keys.

| Genome | Correct patient ID | Correct disease class | Gender | Re-analyzed prioritized gene | Re-analyzed probable diagnostic variant | Mechanisms involved | Max MAF | OMIM diseases | Comments | RegulomeDB score (if less than 4) |
|---|---|---|---|---|---|---|---|---|---|---|
| **007** | X | Eye | F | ZNF644 | 1:91381534:T:G | UTR3 | 0.003 in gnomAD | Myopia 21 | AD inheritance with high myopia | - |
| **007** | X | Eye | F | GRM6 | 5:178413523:G:A, 5:178412718:A:AAAAC | Nonsyn, Intronic indel | 0.002 in gnomAD, Novel | Congenital stationary night blindness | AR inheritance with high myopia | - |
| **009** | W | Eye | F | GPR143 | X:9732545:CACACACACACACA:CG, X:9729836:G:A | Intronic indel, Intronic | Novel, 0.0046 in gnomAD | Ocular albinism with nystagmus | X-linked | - |
| **009** | W | Eye | F | CACNA1F | X:49079668:G:T | Intronic | 0.0038 in gnomAD | X-linked congenital stationary night blindness | Impact mutation by Gerp++, X-linked but didn't find other variant | - |
| **017** | H | Eye | M | ABCA4 | 1:94568686:C:T, 1:94470320:C:T | Nonsyn, Intronic | 0.002 in gnomAD, 0.006 in gnomAD | Retinitis pigmentosa, rod-cone dystropy and other | Nonsyn variant is clinical variant by HGMD (DM?) and other one is impact variant by Gerp++, which is also | - |

| | | | | | | | | | | |
|---|---|---|---|---|---|---|---|---|---|---|
| | | | | | | | | eye disorders | shared by one more patient. But both of these variants are only present in H. | |
| **018** | U | Neurologic | M | CHD2 | 15:93447747:G:C, 15:93446203:T:A | Intronic, Intronic | 0.004 in gnomAD, 0.003 in gnomAD | Epileptic encephalopathy, childhood-onset | Myoclonic encephalopathy, intellectual disability, developmental regression - loss of acquired skills | 2a, - |
| **021** | G | Neurologic | F | KCNMA1 | 10:78792282:G:A, 10:79124696:C:A | Intronic, Intronic | Novel, 0.0003 in gnomAD | Cerebellar atrophy, developmental delay and seizures | Novel intronic variant is impact variant by Gerp++ and other intronic variant is impact variant by CADD and GERP++. | - |
| **030** | R | Neurologic | M | GABRB3 | 15:26792252:G:A, 15:26995302:G:A | UTR3, Intronic | 0.0002 in gnomAD, 0.003 in 1000genome | Childhood absence epilepsy, atonic seizures, intellectual disability | Both variants are impact variants by Gerp++. | - |
| **039** | P | Neurologic | M | GRIN2A | 16:10002180:C:T, | Intronic, Intronic | 0.0025 in gnomAD, 0.0003 | Epilepsy with | Both variants are impact variants by Gerp++ | - |

| | | | | | | | | | | |
|---|---|---|---|---|---|---|---|---|---|---|
| | | | | | 16:9869096: G:A | | in gnomAD | speech disorder | | |
| **042** | O | Eye | F | EYS | 6:66115146: C:T, 6:65312850: G:A | Nonsyn, Intronic | 0.003 in gnomAD , Novel | Retinitis pigmentosa | Retinitis pigmentosa with AR inheritance | - |
| **056** | N | Connective | F | FBN1 | 15:48700642: T:C, 15:48849792: C:A | UTR3, Intronic | 0.00003228 in gnomAD , Novel | Marfan syndrome | We already submitted this variant during challenge time and assessor mentioned it is possible. UTR3 variant is impact variant by GERP++. The novel intronic variant is impact variant by both Gerp++ and CADD. | 3a, - |
| **057** | T | Connective | F | COL5A2 | 2:189897245: T:C | UTR3 | 0.0003 in gnomAD | Ehlers Danlos syndrome | This variant or the other variant is possible. | - |
| **057** | T | Connective | F | COL5A1 | 9:137675325: G:A | Intronic | 0.0012 in gnomAD | Ehlers Danlos syndrome | Both these variants are impact variant by Gerp++. | - |

| | | | | | | | | | | |
|---|---|---|---|---|---|---|---|---|---|---|
| **067** | M | Eye | M | RPGR | X:38138944:CATGTATGAGATAGGCACTTTTGTCCGTGGGTAT:CTTCAATA | Intronic | Novel | X-Linked Retinitis Pigmentosa | Novel Indel in Intronic region | - |
| **067** | M | Eye | M | PDE6B | 4:628493:G:A | Nonsyn | 0.0076 in gnomAD | Retinitis Pigmentosa | This gene is mentioned for RP in pmid: 28912962 | - |
| **068** | J | Neurologic | F | LMNA | 1:156105054:G:T | Nonsyn | Novel | Progeria, muscular dystrophy, lipodystrophy, and others | According to assessor this is possible variant, found by another group. | - |
| **071** | L | Connective | M | FBN1 | 15:48747531:G:C, 15:48807965:T:C | Intronic, Intronic | 0.002 in 1000genome, 0.002 in 1000genome | Joint hypermobility | EDS | - |
| **076** | Q | Connective | F | COL5A1 | 9:137722010:G:GC | FrameShift Insert | Novel | Joint hypermobility | EDS | - |
| **078** | V | Connective | F | COL1A2 | 7:94049587:C:T | Nonsyn | 0.000008 in gnomAD | EDS type III | EDS | - |

| | | | | | | | | | | |
|---|---|---|---|---|---|---|---|---|---|---|
| **078** | V | Connective | F | ARID1B | 6:157264665:G:A | Intronic | 0.0026 in gnomAD | Coffin-Siris Syndrome | Developmental delay, hypoplasia, microcephaly, characteristic facial features | - |
| **079** | K | Connective | M | EZH2 | 7:148538300:A:T | Intronic | Novel | Weaver syndrome | Tall stature, joint hyperflexibility | - |
| **081** | I | Connective | F | TGFB2 | 1:218570954:T:G | Intronic | Novel | Loeys-Dietz syndrome | Connective tissue disorder, pectus excavatum, elongated limbs with joint deformities, inflammatory bowel disease | - |
| **091** | E | Neurologic | F | ARID1B | 6:157465679:C:G | Intronic | Novel | Coffin-Siris Syndrome | Developmental delay with seizures, clinodactyly at 5th finger | 3a |
| **092** | S | Connective | F | ADAMTS2 | 5:178707543:G:A, 5:178726381:TCAGCCGGC:TTAGCCAGG | Intronic, Intronic | 0.000097 in gnomAD, Novel | Ehlers-Danlos syndrome, dermatosparaxix type | AR inheritance | - |

| | | | | | | | | | | |
|---|---|---|---|---|---|---|---|---|---|---|
| **093** | F | Connective | M | COL5A2 | 2:190024916:C:T | Intronic | 0.00003 in gnomAD | EDS | EDS, classical type | - |
| **093** | F | Connective | M | PLEC | 8:145047646:C:T, 8:144989944:C:T | UTR5, UTR3 | Novel, Novel | Epidermolysis bullosa with pyloric atresia | Fragile skin and food intolerance | 2b, - |
| **095** | C | Eye | M | USH2A | 1:215953583:A:G, 1:215964830:T:G | Intronic, Intronic | 0.002 in 1000genome, Novel | Retinitis Pigmentosa | Recessive inheritance | - |
| **097** | D | Connective | F | TNXB | 6:32064371:A:G | Nonsyn | 0.001 in 1000genome | EDS | EDS | 2b |
| **099** | B | Neurologic | M | KCNMA1 | 10:78944523:C:T | Intronic | 0.005 in 1000genome | Paroxysmal nonkinesigenic dyskinesia | Autosomal dominant neurologic disorder with absence seizures, tonic-clonic seizures | - |
| **102** | A | Connective | M | GTF2IRD1 | 7:73973357:C:G | Nonsyn | Novel | William's Syndrome | This gene is important for gene regulation in the brain and in muscles used for movement | - |

| | | | | | | | | | | |
|---|---|---|---|---|---|---|---|---|---|---|
| | | | | | | | | | (skeletal muscles). Characteristic features of Williams syndrome includes the distinctive facial features, dental abnormalities, and problems with visual-spatial tasks such as writing and drawing | |
| **102** | A | Connective | M | COL12A1 | 6:75884801:G:A | Nonsyn | 0.000008 in gnomAD | EDS | EDS - both variants are contributing. | - |

**Supplementary Table S4**: Submitted predictive secondary variants for 24 SickKids5 patients.

| **Genome** | **Predictive Secondary variant** | **dbSNP137** | **Genotype allele depth (ref, alt)** | **Gene** | **Category** | **Frequency** | **Mechanism involved** | **Phenotype description** |
|---|---|---|---|---|---|---|---|---|
| **021** | chr2:48027755:T:C | rs2020912 | 18, 16 | MSH6 | C1 | <= 0.005 | clinical_clinvar_hgmd | Lynch Syndrome |
| **030** | chr7:150644901:G:A | rs199473438 | 28, 24 | KCNH2 | C1 | Novel | clinical_clinvar_hgmd | Romano-Ward Long QT Syndromes Types 1, 2, and 3, Brugada Syndrome |
| **030** | chr2:47707963:TATG:T | - | 9, 11 | MSH2 | C2 | Novel | NonFrameShiftDelete | Lynch Syndrome |
| **030** | chr13:32911607:CCTA:C | rs1064795908 | 17, 19 | BRCA2 | C2 | Novel | NonFrameShiftDelete | Hereditary Breast and Ovarian Cancer |
| **071** | chr2:48027755:T:C | rs2020912 | 14, 16 | MSH6 | C1 | <= 0.005 | clinical_clinvar_hgmd | Lynch Syndrome |
| **076** | chr1:156108510:C:T | rs142000963 | 17, 18 | LMNA | C1 | <= 0.001 | clinical_clinvar_hgmd | Hypertrophic cardiomyopathy, Dilated cardiomyopathy |
| **081** | chr2:47641558:GT:G | rs587779194 | 16, 9 | MSH2 | C1 | Novel | clinical_clinvar_hgmd, Splice Donor | Lynch Syndrome |
| **091** | chr2:47641558:GT:G | rs587779194 | 11, 8 | MSH2 | C1 | Novel | clinical_clinvar_hgmd, Splice Donor | Lynch Syndrome |
